# CTPS1 inhibition synergizes with replication stress signaling inhibition in *MYC*-amplified Group 3 medulloblastoma

**DOI:** 10.1101/2024.06.03.597242

**Authors:** Matthew R Hathaway, Katherine E Gadek, Hawa L Jagana, Isabella C Terrones, John M Hemenway, Aya Miyaki, Ashmitha Rajendran, Michael Meechan, Leonel Elena-Sanchez, Nicholas A Vitanza, Barbara S Slusher, Siobhan S Pattwell, Myron K Evans

**Author notes:** Address correspondence to: Myron K Evans II, Ph.D. Ben Towne Center for Childhood Cancer Research 1920 Terry Avenue Seattle, WA 98101. Financial support: National Cancer Institute of the National Institutes of Health under Award Numbers K22CA262343 (MKE) and K22CA258953 (SSP), Kate Amato Foundation (MKE), Steven Higgins Brain Tumor Research Fund (MKE), Unravel Pediatric Cancer (MKE), the Pediatric Brain Tumor Research Fund of Seattle Children’s (MKE), Seattle Children’s Guild Association, and UW/FHCC/SCH Cancer Center Support Grant (P30CA015704).

## Abstract

*MYC*-driven medulloblastomas (MBs) represent the most aggressive and deadly subgroup of MB, the most common malignant pediatric brain tumor. Direct targeting of MYC itself remains an unmet clinical need, therefore focusing on vulnerabilities driven by MYC may be a viable option for novel therapeutic interventions. Using whole-genome CRISPR screening, we identified the *de novo* pyrimidine synthesis enzyme CTP synthase (*CTPS1*) as a strong dependency in *MYC*-driven MB. CTPS1 is the final and rate-limiting step in the *de novo* pyrimidine synthesis pathway. Targeted inhibition of CTPS1 leads to decreased tumor cell proliferation and markedly reduces MYC expression in G3 MB models. Mechanistically, we demonstrate that single agent CTPS1 inhibition activates the replication stress signaling pathway mediated by ATM-CHK2 and ATR-CHK1. Blockade of CHK1 kinase activity increases sensitivity to CTPS1 inhibition and significantly impedes heterotopic MB tumor growth. CTPS1 enzymatic activity requires the amino acid glutamine, therefore we inhibited CTPS1 using the glutamine antagonists, JHU083 and JHU395. These compounds are prodrugs of 6-diazo-5-oxo-L-norleucine (DON) which were developed to exhibit better tumor targeting and enhanced blood-brain barrier penetrability. Combining JHU083 and CHK1 inhibition demonstrates potent synergy against patient-derived MB xenografts *in vivo*. Our findings strongly suggest that combining *de novo* pyrimidine synthesis and ATR-CHK1 inhibitors is a promising treatment for *MYC*-driven MBs.

**Key Points:**

- *CTPS1* is a unique vulnerability in MYC-driven medulloblastoma
- CTPS1 inhibition activates the ATR-CHK1 replication stress response pathway for cell survival
- Blockade of CTPS1 enzymatic activity synergizes with CHK1 inhibition *in vitro* and *in vivo*

**Importance of the Study:** MYC hyperactivation in tumors drives multiple anabolic processes which contribute to tumor proliferation and aggressiveness in patients. We show that targeting *de novo* pyrimidine synthesis (via CTPS1) limits tumor growth and targets MYC itself through a feedback mechanism. CTPS1 inhibition potently combines with CHK1 blockade and enhances disease control in both heterotopic and orthotopic models of medulloblastoma (MB). Our results support the clinical evaluation of combined CTPS1 and CHK1 inhibition in patients with *MYC*-driven MB.

## Introduction

Despite being the second most common oncological diagnosis, central nervous system tumors are the leading cause of cancer-related mortality in children^1^. Medulloblastoma (MB), the most common malignant pediatric brain tumor, exhibits a complex genomic landscape with diverse molecular subgroups which significantly impact clinical outcomes^2^. Recent advances in genomic, epigenomic, and proteomic profiling have identified four subgroups of MB: WNT, sonic hedgehog (SHH), Group 3, and Group 4. Patients with Group 3 MB account for ∼25-30% of all diagnosed cases and have the poorest overall survival of all groups^2^.

Dysregulation of the proto-oncogene MYC is a hallmark of G3 MB, with patients exhibiting recurrent gene amplification and genomic fusions^3^ or high MYC activity due to increased protein stability^4^. Despite our understanding of MYC as a critical oncogene^5^, no direct targeted agents have been tested clinically in pediatric patients. This has motivated the study of synthetic lethality approaches, targeting vulnerabilities specifically driven by MYC hyperactivation^6^.

MYC supports cancer cell proliferation through numerous mechanisms, including rewiring of cellular metabolism, by modulating anabolic processes to sustain the heightened demands of rapidly dividing cells. Oncogenic MYC drives the upregulation of key enzymes crucial to nucleotide biosynthesis, ensuring a robust supply of purine and pyrimidine precursors needed for rapid tumor cell proliferation^7^. Simultaneously, MYC promotes metabolic rewiring of glycolysis and the pentose phosphate pathway (PPP), channeling resources toward nucleotide synthesis^7^. This dynamic interplay is a characteristic feature of G3 MB^8^. Further, it underscores the significance of MYC in tailoring cellular metabolism to meet the unique demands of oncogenesis, unveiling potential vulnerabilities that could be harnessed for targeted therapeutic interventions. Inhibition of nucleotide synthesis and metabolism effectively induces cell death and is commonly exploited by various chemotherapeutics sometimes used in the treatment of MB^9,10^. However, these treatments commonly yield unwanted side effects in patients, suggesting that targeting tumor specific metabolism pathways could alleviate negative sequelae. CTP synthase 1 (CTPS1), the rate limiting enzyme in the *de novo* synthesis of the nucleotide CTP, is one such metabolic target^11,12^. CTPS1 has previously been shown as a vulnerability in various adult solid and hematologic tumors^13–16^ and a small molecule inhibitor of CTPS1 is currently in clinical trials (NCT05463263). Further, a recent paper linked MYC expression to CTPS1 inhibitor sensitivity, demonstrating synthetic lethality in colorectal cancer^17^. Low on-target, off-tumor toxicity of this molecule (and similar compounds) is most likely due to significant compensation by the homolog CTPS2 in most tissues, excluding the lymphoid compartment post-infection^18,19^.

In this study, using functional genomics and small molecule inhibition, we characterize CTPS1 as a targetable vulnerability in G3 MB and identify MYC-amplified MB as exquisitely dependent on *de novo* pyrimidine but not purine synthesis. Unlike in hematological malignancies where CTPS1 inhibition alone significantly inhibits growth and increases apoptosis^13,16^, we observe limited cytotoxicity as a single agent in G3 MB. We demonstrate that inhibition of CTPS1 in G3 MB activates the ATR-CHK1 replication stress response, enhancing cell survival; dual antagonism of *de novo* pyrimidine synthesis and CHK1 synergizes *in vitro* and *in vivo*. Finally, we demonstrate that a blood-brain barrier (BBB) penetrant inhibitor of nucleotide biosynthesis combines with CHK1 inhibition increasing survival in an orthotopic model of MB. Our findings identify CTPS1 as a novel target and demonstrate that this combinatorial approach may be a valid option to move forward clinically for patients with G3 MB.

## Results

### CTPS1 and the *de novo* pyrimidine synthesis pathway are dependencies in Group 3 medulloblastoma

Functional genomics screens allow for unbiased identification of potentially druggable vulnerabilities in tumors and have been utilized to guide pre-clinical work and clinical trials in both adult and pediatric oncology. To evaluate dependencies specific to Group 3 medulloblastoma (MB), we first explored the Pediatric Cancer Dependency Map (DepMap)^20^. Our analysis of 4 well-characterized G3 MB cell lines (D283, D341, D425, and D458) versus all other pediatric entities, revealed CTP synthase 1 (*CTPS1*) as the third-strongest dependency (defined by a negative CHRONOS score, **Figure 1A**). Comparing G3 MB to only pediatric (peds) central nervous system (CNS) tumors still reveals a strong dependency on *CTPS1* (**Figure S1A**), suggesting a specific function in *MYC*-amplified MB. CTPS1 is the rate limiting enzyme responsible for the glutamine-dependent conversion of uridine triphosphate (UTP) to cytidine triphosphate (CTP), an essential step of the *de novo* pyrimidine synthesis pathway^12^ (**Figure 1B**). Further analysis revealed significant dependencies on other enzymes in the *de novo* pyrimidine synthesis pathway including carbamoyl-phosphate synthetase 2, aspartate transcarbamylase, and dihydroorotase (*CAD*), uridine monophosphate synthase (UMPS), and dihydroorotate dehydrogenase (*DHODH*) (**Figure 1C**, G3 MB vs. all peds, **S1B** G3MB vs. peds CNS). UMP-CMP kinase (*CMPK1*) scores as a vulnerability however, no difference is seen when compared to other pediatric tumors. This is further validated by a recent publication which revealed DHODH as a druggable target by CRISPR screening in another G3 MB cell line (SUMB002)^21^. Mining of CRISPR screening data in two fetal neural stem cell (NSC) lines^22^ shows no dependency on *CTPS1* or other *de novo* pyrimidine synthesis enzyme genes (**Figure S1C**). Interestingly, neither disruption of *CTPS2* (**Figure 1C, S1B**) or genes in the salvage pathway (**Figure S1D**) had any effect on cell growth, suggesting a strong, non-redundant role for *CTPS1* in G3 MB.

**Figure 1:**
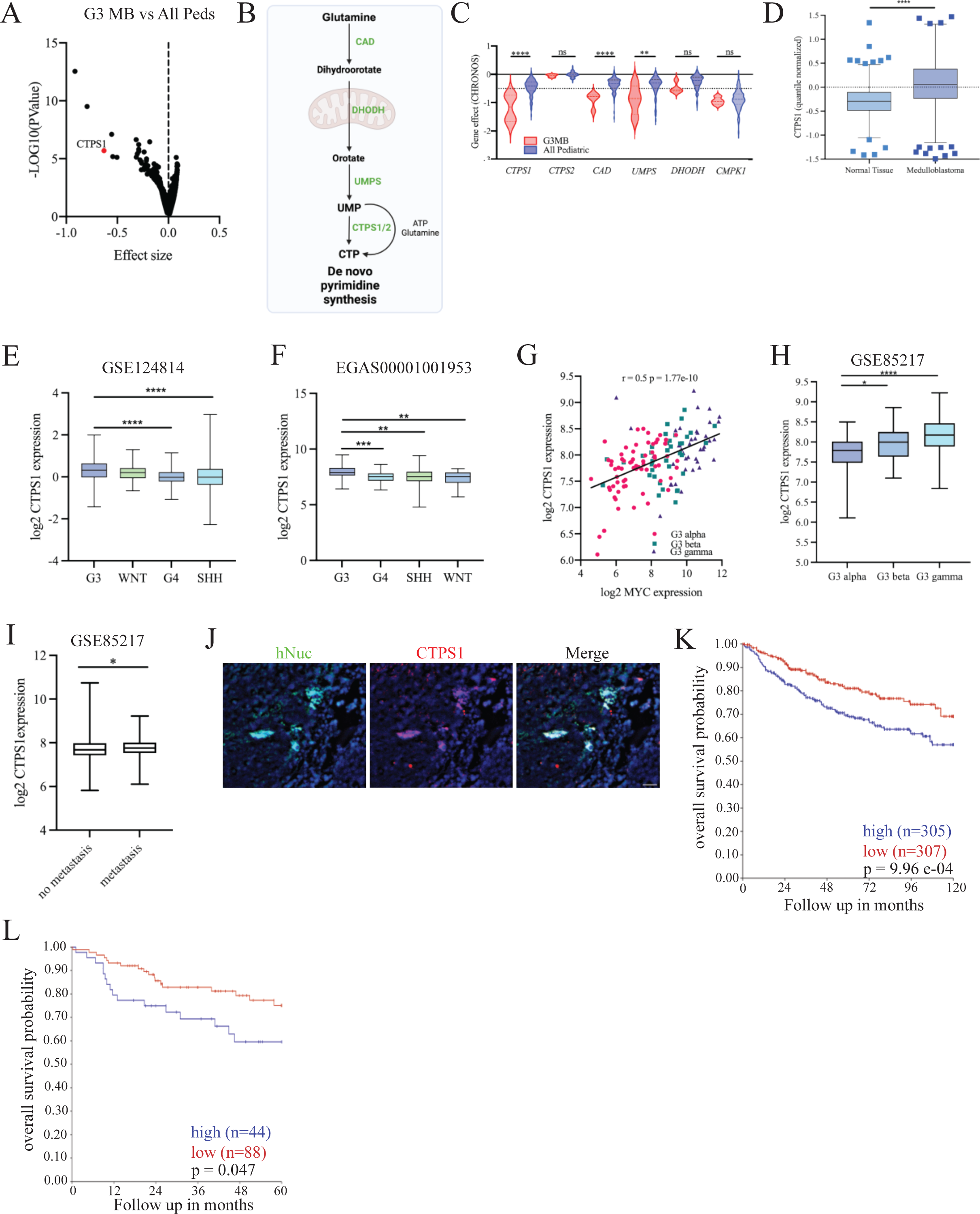
*CTPS1* is overexpressed in medulloblastoma and is a dependency in G3 MB. **a,** Volcano plot comparing CERES scores of G3 MB cell lines (D283, D425, D458, D341) versus other pediatric tumor lines (n=111) in the Dependency Map using two class comparison. **b,** Diagram of *de novo* pyrimidine synthesis pathway, **c,** CERES gene dependency scores for pyrimidine synthesis genes in G3 MB cells (n=4) compared to other pediatric cell lines (n=111) in the DepMap, 23Q4 release. **d,** Quantile normalized expression of CTPS1 in normal (n=291) or primary medulloblastoma (n=1350) samples. **e-f,** Log_2_ normalized expression of CTPS1 in GSEs (e-GSE124814, f-EGAS00001001953). Samples are separated by MB subgroup. **g,** Linear regression of log_2_ normalized CTPS1 and MYC expression from GSE85217 separated by subtype. **h,** Log_2_ normalized expression of CTPS1 in G3 MB subtypes from same samples in **g**. **i,** Log_2_ normalized CTPS1 expression in primary tumor samples from patients with or without metastases at diagnosis from GSE85217. **j,** Immunohistochemistry of G3 patient-derived xenograft tissue after orthotopic implantation in nude mice (red-CTPS1, green-human nuclear antigen (hNuc), blue-DNA). Scale bar=100 μM. **k,** Kaplan-Meier survival of patients diagnosed with MB separated by median expression of *CTPS1*, GSE85217. **l,** Kaplan-Meier survival of *Myc*-high G3 MB patients separated by top 1/3 versus bottom 2/3 expression of *CTPS1*. For **k** and **l**, number of patients are shown on inset. *, P < 0.05; **, P < 0.01; ***, P < 0.001, Data are means ± SEM.

### Higher *CTPS1* expression is associated with disease advancement and poor prognosis in MB

Using a batch-normalized bulk RNA-seq resource (GSE124814)^23^ comprised of 1350 MB samples and 291 normal cerebellum revealed increased *CTPS1* expression in tumors compared to normal cerebella (**Figure 1D**). Subgroup analysis uncovered the highest average expression of *CTPS1* in G3 tumors in two independent datasets (p<0.05 G3vs.G4 and G3vs.SHH, **Figure 1E-F**). Analysis of a third dataset (GSE85217)^24^ showed a strong correlation between *MYC* and *CTPS1* expression in G3 MB tumors, with the highest expression in those tumors characterized as Group3γ (**Figure 1G-H)**. *CTPS1* expression also correlates with metastatic status of patients at time of diagnosis (**Figure 1I**), further demonstrating association with aggressive characteristics. We validated these RNA-seq findings in a complementary proteomic dataset^4^. In this data, G3 MB tumors were subclassified as G3a and G3b, where G3a tumors are characterized by a MYC-activated state either by gene amplification or post-translational alterations which increase MYC protein stability. We demonstrate higher CTPS1 protein levels in G3a vs. G3b tumors (**Figure S1E**), as well as higher expression of other components of the *de novo* pyrimidine pathway (**Figure S1F**). Immunohistochemistry of G3 MB patient-derived xenografts confirms expression of CTPS1 in tumor tissue with little expression in normal mouse tissue (**Figure 1J**). Weaker or no correlations were seen with other genes in the pyrimidine synthesis pathway (**Figure S1G**). A correlation between *CTPS1* and *CCNB1*, a key regulator of cell cycle progression and a proposed risk stratification gene^25^, was also noted (**Figure S1H**). Consistent with this, higher *CTPS1* expression was associated with worse overall survival in all MB patients (**Figure 1K**) and in G3 MB patients with high MYC expression (**Figure 1L**). Of the other *de novo* pyrimidine synthesis pathway enzymes, only *DHODH* and *UMPS* expression correlated with poor survival in G3 MB patients (**Figure S1I**).

### Targeting CTPS1 reduces MB cell proliferation *in vitro*

Based on the significant attenuation of growth using CRISPR-Cas9 targeting of *CTPS1*, we targeted *CTPS1* using shRNA. After selection with puromycin, sphere formation was assessed and cell proliferation measured by BrdU incorporation. In two different G3 MB cell lines, cells exhibited decreased sphere forming capacity (**Figure 2A-B**), and diminished proliferation concurrent with downregulation of *CTPS1* expression (**Figure 2C**). Despite significant growth inhibition *in vitro*, animals implanted with sh*CTPS1* cells showed limited survival benefit compared to control shRNA implanted animals. HD-MB03 sh*CTPS1* tumors retained CTPS1 expression suggesting a negative selection for cells with *CTPS1* knockdown *in vivo*. We next employed treatment with 3-Deazauridine (DAU) which is a structural analog of uridine that competitively inhibits the synthesis of CTP by both CTPS1 and CTPS2^26^. DAU treatment caused a dose-dependent decrease in cell proliferation across a panel of G3 MB cell lines (**Figure 2D**). This inhibition could be reversed by supplementation of either uridine or cytidine, but not adenosine or guanosine (**Figure 2E**). Similar to previously published data^14,27^, CTPS1 pharmacological inhibition resulted in S-phase blockade with decreased cells in G_0_-G_1_, which was reversed by addition of cytidine or uridine (**Figure 2F, S2A**). Furthermore, DAU treatment also decreased MYC expression in tumor lines similar to previous reports with DHODH inhibition in G3 MB^21^ (**Figure 2G**).

**Figure 2:**
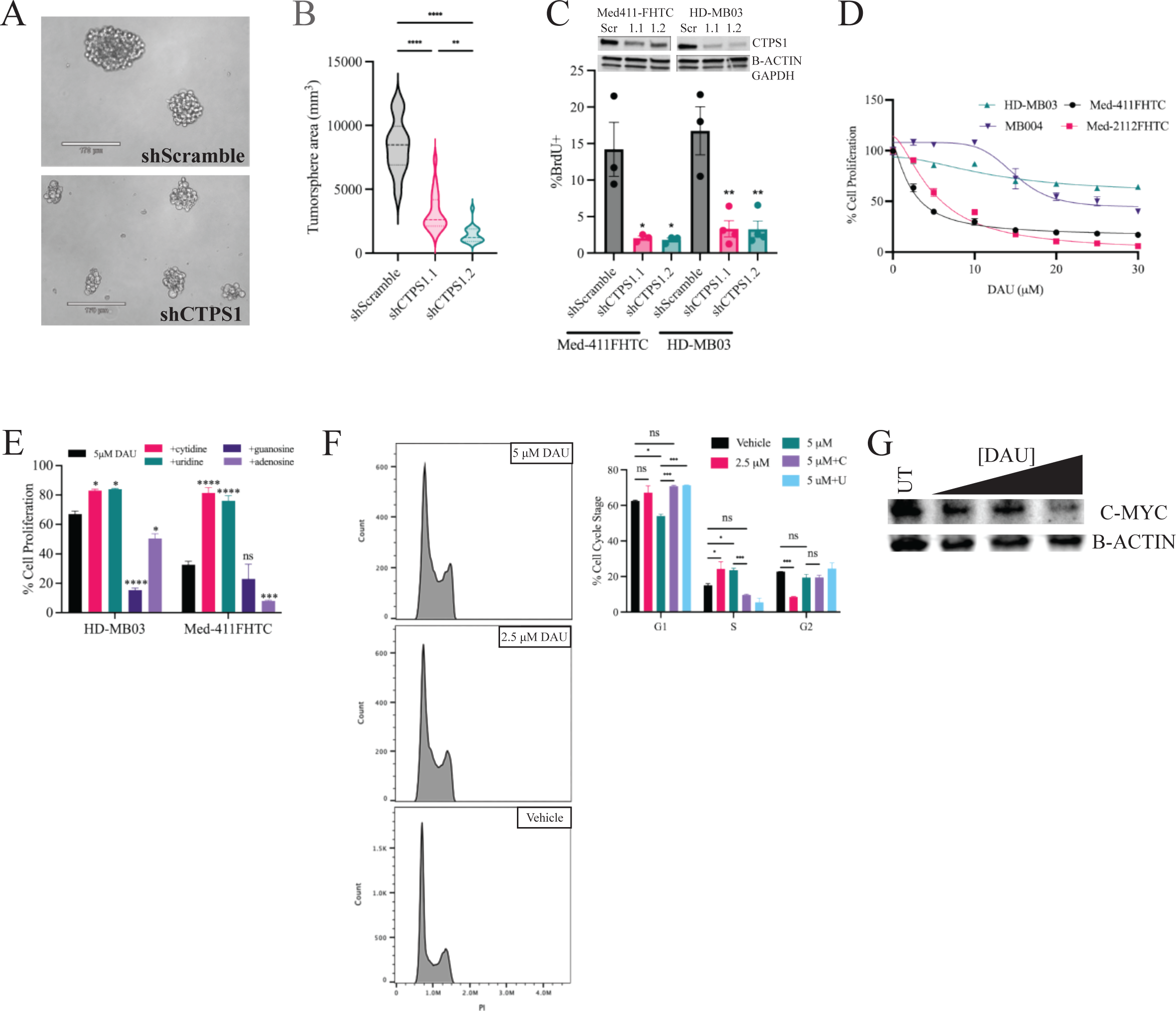
CTPS inhibition suppresses cell proliferation in G3 MB tumor cells. **a,** Representative images of HD-MB03 tumorspheres transduced with indicated shRNAs 72h after plating. **b,** Quantification of area of tumorspheres formed in shScramble, shCTPS1.1, and shCTPS1.2 transduced HD-MB03 cells. **c,** Quantification of FACS for BrdU-positive HD-MB03 and MED-411FHTC cells transduced with indicated shRNAs (n=3). *Inset*, Western immunoblot analysis of CTPS1 expression in transduced cells in **a-c** Scr= scramble, 1.1 =shCTPS1.1, 1.2= shCTPS1.2. **d,** Cell proliferation analysis of HD-MB03, MB004, Med-411FHTC, and Med-2112FHTC tumorspheres treated with indicated concentrations of DAU for 72h. **e,** Cell proliferation analysis of HD-MB03 and Med-411FHTC tumorspheres treated with vehicle or DAU (5μM) alone or in the presence of 50μM cytidine, uridine, adenosine, or guanine (n=3). **f,** Cell-cycle analysis of HD-MB03 cells treated with the indicated concentrations of DAU alone or in the presence of cytidine (C) or uridine (U) for 48h (n=3). **g,** Western immunoblot analysis of MYC expression in HD-MB03 cells treated with increasing doses of DAU. *, P < 0.05; **, P < 0.01; ***, P < 0.001, Data are means ± SEM.

### G3 MB cells are dependent on *de novo* pyrimidine but not *de novo* purine synthesis

Previous work in other tumor types has revealed that MYC regulates both pyrimidine and purine biosynthesis through direct regulation of multiple enzyme-encoding genes in both pathways^7^. Analysis of previously published ChIP-seq data from two human G3 MB tumors demonstrates significant MYC binding at the *CTPS1* promoter, with lower binding at the *CTPS2* promoter correlating with gene expression^28^ (**Figure S3A,B**). Analysis of the same dataset from **Fig 1D**. revealed strong correlation between *MYC* and numerous genes encoding enzymes in the *de novo* purine synthesis pathway (**Figure S3C**). Exploration of DepMap data revealed that while some genes in the pathway scored as dependencies, they were no different than pediatric CNS tumors (**Figure S3D**). The only purine related enzyme that scores is thymidylate synthetase (TYMS), a catabolic one-carbon metabolism enzyme that has an indirect effect on purine by driving thymidine synthesis from dUMP salvage (**Figure S3E**). This correlates with activity of TYMS inhibitors pemetrexed and 5-fluorouracil^29,30^ and suggests the possibility of recently developed novel inhibitors for future pre-clinical modeling^31,32^.

To further clarify that inhibition of pyrimidine but not purine synthesis was antagonistic to growth of G3 MB cells, we treated both Med-2112FHTC and Med-411FHTC with lometrexol (LTX), an inhibitor of the purine biosynthetic enzyme glycinamide ribonucleotide formyltransferase (GARFT). We observed no effect on cell fitness after treatment with LTX, similar to previous reports in IDH-mutant glioma and pediatric diffuse midline glioma^33^ (**Figure S3F**).

### CTPS1 inhibition alone is cytostatic but combines with CHK1 inhibition

Western blot analysis of cells treated with DAU revealed little change in PARP cleavage, a classic marker of apoptosis, at doses up to 5μM suggesting a cytostatic mechanism rather than cytotoxicity (**Figure 3A**). Diehl et.al. recently published that nucleotide imbalance, caused by targeting pyrimidine and/or purine enzymes, activates replication stress signaling sensed by the ATR and ATM kinases^27^. These kinases sense both single- and double-stranded breaks and phosphorylate CHK1 and CHK2, respectively, which mediate survival and recovery of cells. We observed increased phosphorylation of both CHK1 and CHK2 after treatment with DAU (**Figure 3B**). We then combined targeting of CTPS1 with the potent CHK1/2 inhibitor prexasertib, which previously showed efficacy in G3 MB cells^34^. Combining sublethal doses of prexasertib (**Figure S4A**) led to a dose-dependent decrease in the IC_50_ of DAU in both HD-MB03 and Med-411FHTC cells (**Figure 3C and D**). **Table 1** lists synergy scores as calculated by multiple reference models^35^ which revealed potent synergy between CTPS1 and CHK1/2 inhibition in G3 MB. This combination also led to apoptosis (**Figure 3C, inset** – cleaved PARP) and significant DNA damage (measured by pH2AX) (**Figure 3E-F**). Prexasertib treatment also synergized with BRQ, suggesting that CHK1 inhibition may broadly combine with antagonism of *de novo* pyrimidine synthesis in G3 MB (**Figure S4B**). Utilizing a heterotopic xenograft model of G3 MB in athymic nude mice, we assessed the efficacy of combining DAU and prexasertib on tumor growth (**Figure 3G**). While single agent targeting of either CTPS1 or CHK1 led to delays in tumor growth, the combination of DAU and prexasertib synergized and completely abrogated growth in this model (**Figure 3H**). Immunohistochemical analysis revealed decreased Ki67 corroborating our doubling time findings (**Figure 3I**). Phosphorylation of H2AX was also elevated in combination treated animals. Western blots revealed significant cleavage of PARP in our combination treatment and sustained phosphorylation of CHK1/CHK2 (S345/S516) in combination treatments (**Figure 3J**). We also noted decreased phosphorylation of the CHK1 target CDC2 (Y15) in some combination treatment mice as well.

**Figure 3:**
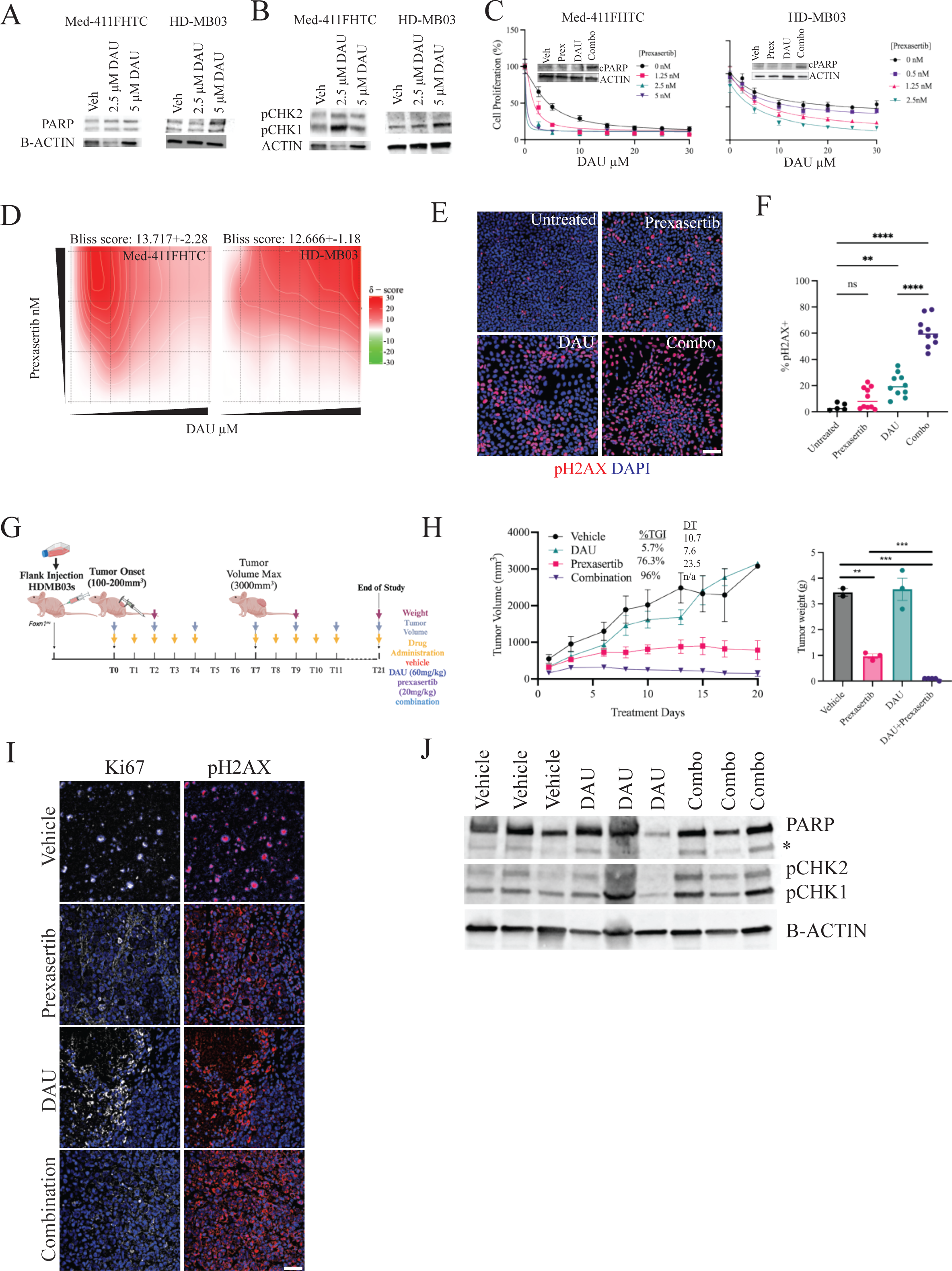
CTPS1 inhibition synergizes with ATR-CHK1 blockade *in vitro* and *in vivo*. Western immunoblot for **a,** PARP and **b,** phospho-CHK1/2 in Med-411FHTC or HD-MB03 cells treated with vehicle, 2.5μM, or 5μM DAU for 48h. **c,** Cell proliferation analysis of Med-411FHTC (left) and HD-MB03 (right) tumorspheres with increasing concentrations of DAU alone or in the presence of indicated prexasertib (CHK1 inhibitor) concentrations (n=3). *Inset*, Western immunoblot analysis of cleaved PARP in cells treated with prexasertib alone (1.25nM), DAU alone (2.5μM), or the combination. **d,** 2D contour plots of synergy (Bliss calculation) between DAU and prexasertib in Med-411FHTC (left) and HD-MB03 (right) tumorspheres calculated by Synergy Finder. **e,** Immunofluorescence for phospho-H2AX in D283 G3 MB cells treated as indicated. Scale bar 25 μM. **f,** Quantification of images in **e** (n=5 images per condition, experiment repeated twice). **g,** Pre-clinical mouse treatment protocol for data shown in **h**. **h,** left – Tumor volume of HD-MB03 flank xenografts treated as indicated n=5 (vehicle) or 6 (all others). Tumor volume statistics (p-value) – veh-prex: 3.9E-6, veh-combo: 3.1E-11, prex-combo: 8.7E-3. TGI: tumor growth inhibition DT: doubling time; right – Quantification of tumor weights in grams at Day 20 post-treatment (veh, n=2, DAU, n=3, prex, n=3, combo, n=5). **i,** Representative images of immunohistochemistry for Ki-67 and phospho-H2AX in tumors at endpoint. **j,** Western immunoblot for total PARP (* denotes cleaved PARP) and phospho-CHK1/2 in tumors from **h,**. *, P < 0.05; **, P < 0.01; ***, P < 0.001, Data are means ± SEM.

**Table 1:** Synergy scores for CTPS1 inhibitiors with prexasertib.

|  | Drug 1 | Drug 2 | ZIP_synergy | ZIP_synergy_p_value | HSA_synergy | HSA_synergy_p_value | Loewe_synergy | Loewe_synergy_p_value | Bliss_synergy | Bliss_synergy_p_value |
| --- | --- | --- | --- | --- | --- | --- | --- | --- | --- | --- |
| HD-MB03 | prexasertib | DAU | 12.60639662 | 5.33E-20 | 17.40683667 | 3E-117 | 10.83592936 | 0.000055 | 12.0364247 | 6.25E-15 |
|  | prexasertib | JHU395 | 5.489365734 | 0.000469 | 8.794607635 | 2.34E-08 | -1.769786284 | 0.886 | 5.237058489 | 0.00347 |
|  | prexasertib | JHU083 | 6.122734406 | 0.0211 | 12.55551008 | 8.69E-21 | 9.364874489 | 0.000188 | 5.965281089 | 0.027 |
| Med-411FHTC | prexasertib | DAU | 11.03533493 | 4.03E-19 | 13.89670691 | 2.03E-92 | 8.049172603 | 0.101 | 10.78442356 | 2.35E-14 |
|  | prexasertib | JHU395 | -1.533340467 | 0.655 | -4.356167358 | 0.0353 | -7.164625504 | 0.172 | -3.25086577 | 0.396 |
|  | prexasertib | JHU083 | 8.805722263 | 0.0000897 | 18.33827545 | 6.16E-18 | 4.763895713 | 0.534 | 8.852162691 | 0.0000665 |

### Targeted glutamine antagonism combines with CHK1 inhibition in orthotopic models

While our *in vitro* and *in vivo* data with DAU are striking, there is only one reference to blood-brain barrier penetration of this compound in humans^36^. This was evaluated in a handful of patients with intracerebral tumors which commonly have abnormally leaky vasculature^37^. MBs, apart from the WNT subgroup, are characterized to have a fully intact BBB which modulates response to BBB-impermeable drugs^38^. The combination of these make DAU an unlikely candidate for G3 MB therapeutic development. CTPS enzymatic activity requires the amino acid glutamine as well as ATP to produce CTP^39^. Glutamine is also required at the first step of the *de novo* synthesis pathway catalyzed by CAD (**Figure 1B**). Interestingly, the glutamine antagonist 6-diazo-5-oxo-l-norleucine (DON) was previously shown to be an irreversible inhibitor of multiple nucleotide synthesis enzymes^40^, including CTPS1 and CAD. Single agent DON administration leads to significant phosphorylation of CHK1/2 (**Figure 4A**) and cell death in G3 MB tumor cells^41^. We utilized two prodrugs of DON – 1) isopropyl 6-diazo-5-oxo-2-(((phenyl (pivaloyloxy) methoxy) - carbonyl) amino) hexanoate (JHU395) and 2) Ethyl 2-(2-Amino-4-methylpentanamido)-DON (JHU083) – which exhibit enhanced lipophilicity, brain penetration, and were previously shown to target G3 MB tumor cells^41,42^ (**Figure S5A**). We first tested JHU083 and JHU395 as single agents in our patient-derived cell lines which revealed efficacy with inter-cell line variability (**Figure 4B**). Co-administration of prexasertib synergized with both JHU083 (**Figure 4C and D**) and JHU395 (**Figure 4E and F**), decreasing the IC_50_ in multiple G3 MB tumor lines. Synergy analyses (**Table 1**) revealed stronger synergism of prexasertib with JHU083 than with JHU395, so for the remainder of our study we used the combination of JHU083+prexasertib. Combining JHU083 and prexasertib led to significant accumulation of DNA damage (**Figure 4G-H**) similar to DAU+prexasertib. Previous work demonstrated significant BBB penetration and tumor accumulation of both JHU083^42^ and prexasertib^43^ in G3 MB tumor bearing mice, therefore we tested single agents and combinations after orthotopic implantation of luciferase-expressing G3 MB cells. After confirmation of tumor growth by IVIS imaging, mice were randomized to one of four groups and treated with JHU083 alone or in combination with prexasertib (**Figure 4I**). While single agent administration had limited effect on tumor growth or animal survival, the combination of JHU083 and prexasertib significantly extended the median survival in animals implanted with either HD-MB03 (**Figure 4J, S5B**) or Med-411FHTC (**Figure 4K, S5C**). Further, this combination as well as our DAU+prexasertib combination had limited systemic toxicities as measured by weight loss in animals (**Figure S5D,E**). Western immunoblot analysis of tumors revealed increased phospho-CHK1/2 and phospho-CDC2 in JHU083 single agent treated tumors. Combination treatment led to a blockade of CHK1/2 and CDC2 phosphorylation as well as a decrease in proliferation as measured by PCNA (**Figure 4L**).

**Figure 4:**
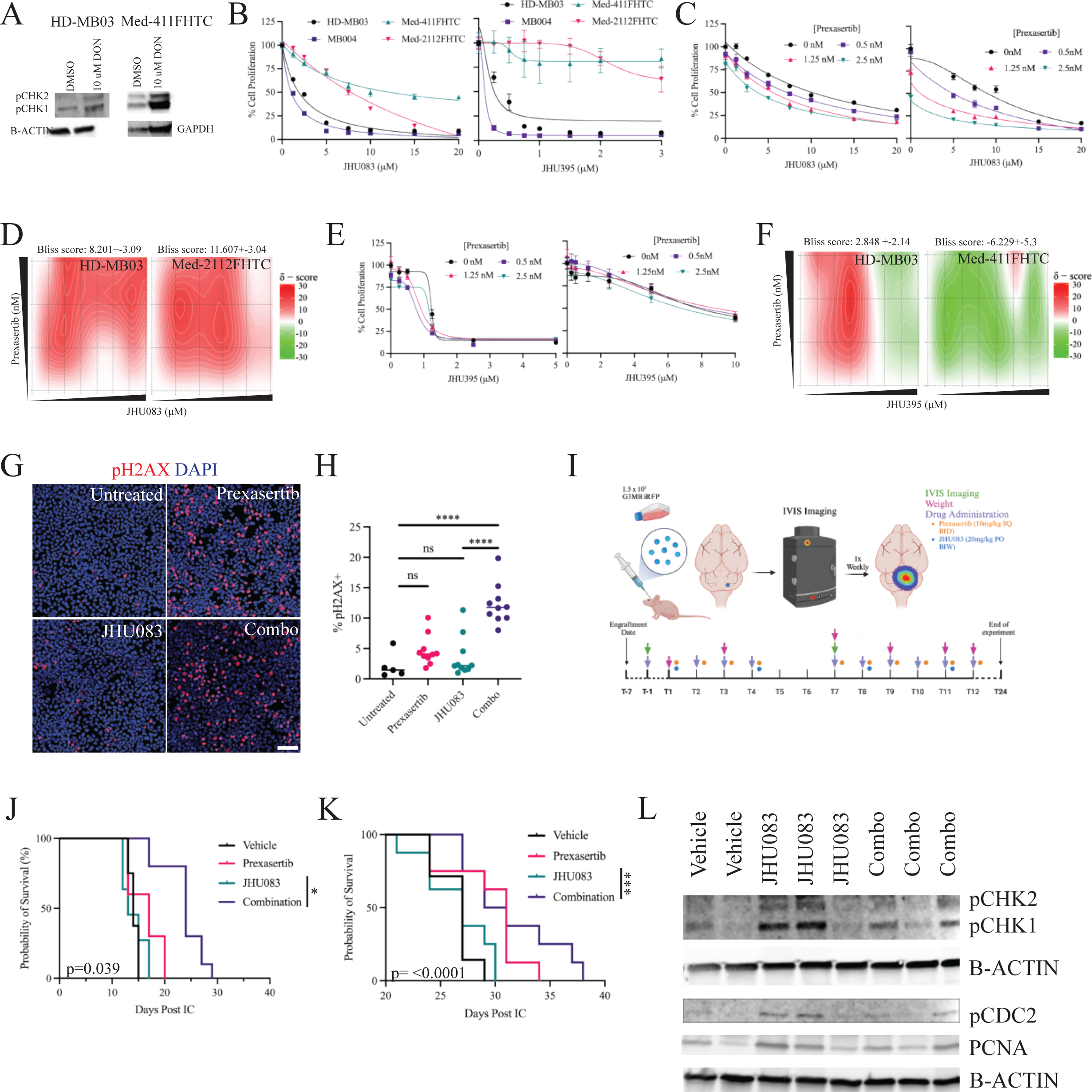
CTPS1 inhibition via glutamine analogs synergizes with CHK1 inhibition in orthotopic G3 MB models. **a,** Western immunoblot for phospho-CHK1/2 in Med-411FHTC (left) and HD-MB03 (right) tumorspheres treated with 10μM DON. **b,** Cell proliferation analysis of HD-MB03, MB004, Med-411FHTC, and Med-2112FHTC tumorspheres treated with indicated concentrations of JHU083 (left) and JHU395 (right) for 72h (n=3). Cell proliferation analysis and 2D contour plots of synergy (Bliss calculation) for **c, d** JHU083 and **e, f** JHU395 alone or in the presence of indicated prexasertib concentrations (n=3). **g,** Immunofluorescence for phospho-H2AX in D283 G3 MB cells treated as indicated. Scale bar 25 μM. **h,** Quantification of images in **g** (n=5 images per condition, experiment repeated twice). **i,** Pre-clinical mouse treatment protocol for data shown in **j-k**. Kaplan-Meier survival of mice orthotopically implanted with **j,** HD-MB03 or **k,** Med-411FHTC cells transduced with luciferase expressing construct and treated as indicated. **l,** Western immunoblot for phospho-CHK1/2, phospho-CDC2, and PCNA in tumors from **j-k**. *, P < 0.05; **, P < 0.01; ***, P < 0.001, Data are means ± SEM.

## Discussion

Herein, we report a dependency on *CTPS1* in *MYC*-amplified MBs when compared to other pediatric malignancies. Interrogation of patient data revealed high expression of *CTPS1* in tumors compared to normal brain (cerebellum/upper rhombic lip) with *MYC*-amplified G3γ tumors displaying the highest expression. This was confirmed using recently published proteomics data which correlates CTPS1 with G3a tumors, characterized by increased MYC expression and activity^4^. Consistent with these findings, *CTPS1* expression correlates with poor survival in *MYC* high patients. CTPS1, along with its homolog CTPS2, catalyze the rate limiting step of the *de novo* pyrimidine biosynthesis pathway. Interestingly, *MYC*-amplified MB tumors do not show a dependency on CTPS2, which also shows no correlation with survival. This is consistent with previous reports that demonstrate differential activity of the two enzymes, despite ∼70% protein sequence homology^44^. Further analysis revealed vulnerabilities to deletion of other members of the *de novo* pyrimidine biosynthesis pathway, confirming previous reports of altered pyrimidine metabolism in MYC-amplified MB^21^, and more broadly across a number of other brain tumor types^33^. Furthermore, recent work in a chemotherapy-resistant model of MB revealed upregulation of pyrimidine metabolic processes in resistant tumors^45^, suggesting utility of CTPS1 targeting in both upfront and recurrence settings.

Using shRNAs and a small molecule inhibitor of CTPS1 (3-deazuridine, DAU) we demonstrate that single agent targeting of CTPS1 leads to cell cycle arrest and decreased proliferation which can be rescued by addition of exogenous pyrimidine nucleosides *in vitro*. This targeting of CTPS1 also leads to a decrease in MYC expression which may further impact tumor growth. Interestingly, germline mutations in *CTPS1* lead to no overt phenotypes outside of the hematopoietic system, suggesting that inhibition may deliver potent on-target effects in tumors while limiting deleterious side effects in patients^18,19^. This is highlighted by single agent efficacy of DAU in our flank model with no significant weight loss observed. Mechanistically, amplified or hyperactive MYC in G3 MB induces expression of CTPS1 required to synthesize nucleosides to maintain replicative capacity of tumors. Inhibition of CTPS1 limits the pool of CTP available for replication^9^, leading to activation of the replication stress response pathway mediated by ATR-CHK1/ATM-CHK2 and subsequent cell cycle and proliferation arrest. Combined targeting of CTPS1 and the replication stress pathway forces cells to continue through the cell cycle, accumulating significant DNA damage and undergoing apoptosis (**Figure 5**).

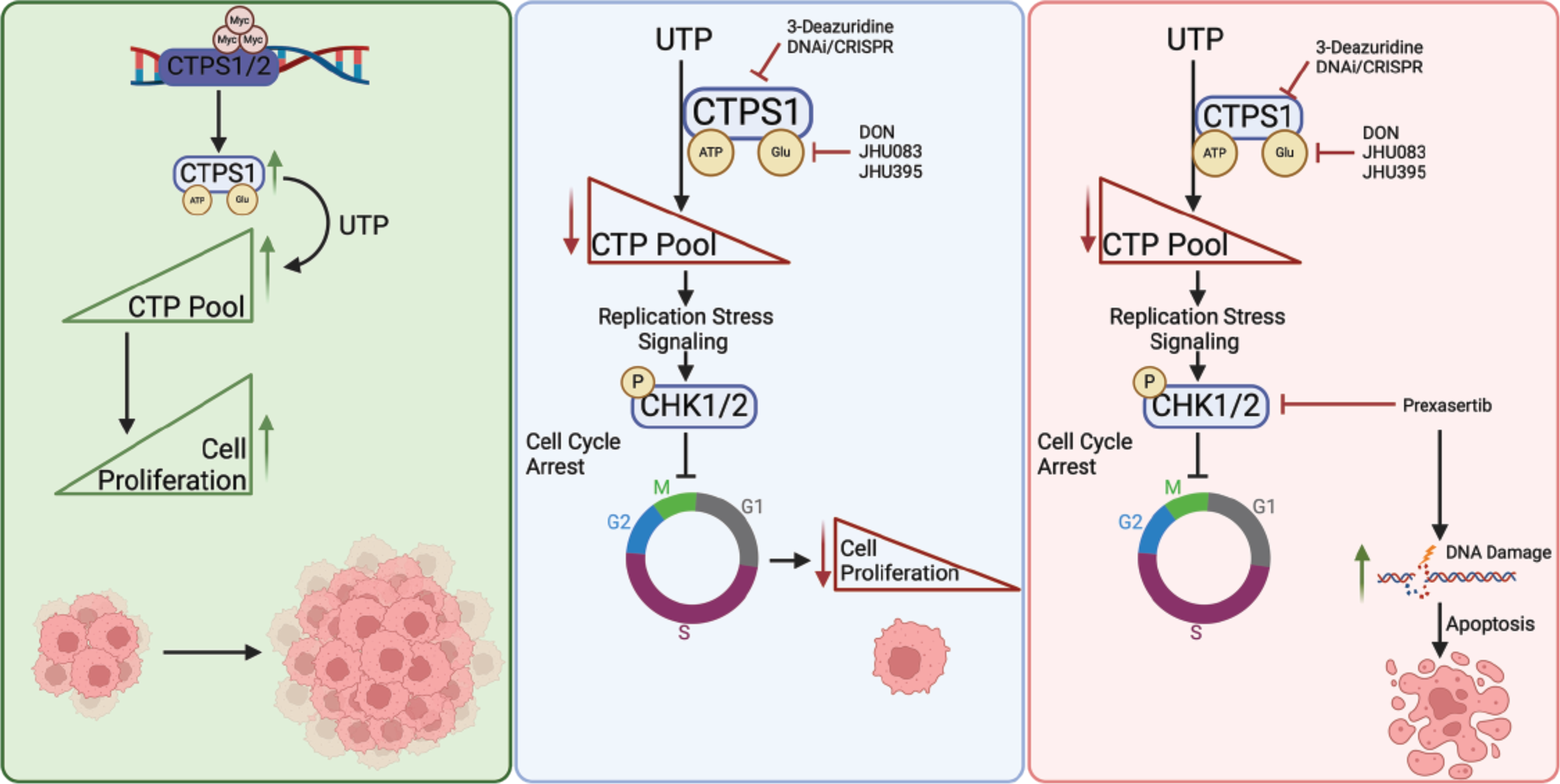

This phenomenon has been observed in other tumor and normal cell types wherein combining nucleotide depletion with ATR/ATM inhibitors leads to cell death^14,17,27^. We demonstrate significant synergy between the CHK1 inhibitor, prexasertib, and DAU both *in vitro* and *in vivo*. Further, we also demonstrate that the DHODH inhibitor, brequinar, also synergizes with prexasertib suggesting the utility of dual *de novo* pyrimidine synthesis and replication stress signaling inhibition in treatment of *MYC*-amplified MB. Identification of this combination may further enhance the clinical utility of a DHODH inhibitor from Bayer which shows potent BBB penetration, but whose clinical development was halted due to single agent inactivity (NCT03404726).

Excitingly, CTPS1 inhibitors have recently entered clinical development (NCT05463263) for hematological malignancies; however, blood-brain barrier (BBB) penetrance has not been assessed. This compound (STP938) mimics CTP and acts an allosteric inhibitor of enzymatic activity; impeding CTPS1 binding to UTP. CTPS1 also has a glutamine amidotransferase domain which hydrolyzes glutamine to ammonia which is required for catalysis of 4-phosphoryl UTP to CTP. Previous work identified that the glutamine antagonist 6-Diazo-5-oxo-L-norleucine (DON) binds to this domain, as well as other glutamine utilizing enzymes, to inhibit covalent binding of glutamine^40^. We observed a similar activation of CHK1/2 when cells were treated with DON as with DAU. While DON does accumulate in mouse models of brain tumors, previous clinical trials were unsuccessful due to dose-limiting toxicities^46^. We utilized two DON prodrugs, JHU083 and JHU395, with enhanced BBB penetrability and tumor-targeting, leading to reduced toxic side effects^47,48^. These inhibitors combine strongly with CHK1 inhibition *in vitro*, with the combination of JHU083 and prexasertib showing significant synergy. We tested this combination *in vivo* and noted increased survival, decreased proliferation, and increased apoptosis, and limited weight loss despite prolonged combination treatment. While our studies are done in immune-deficient models, recent studies have identified immune modulatory activity of JHU083^49,50^ which may bolster effects of this drug in immune-competent mouse models and patients. Together, the data presented herein identify a targeted, combinatorial approach for the treatment of *MYC*-amplified MB patients who currently have the least favorable outcomes.

## Methods

### Cell culture

Patient derived medulloblastoma cell lines (Med-411FHTC and Med-2112FHTC, a gift from J. Olson) were grown in NeuroCult NS-A Basal medium (Stemcell Technologies, 05750) with Neurocult NS-A Proliferation Supplement (Stemcell Technologies, 05753), and 1% Penicillin/Streptomycin (Sigma-Aldrich, P4333). HD-MB03 (obtained from T. Milde) and MB004 (a gift from P. Bandopadhayay) cell lines were grown in Gibco Neurobasal medium (Gibco, 21-103-049) with 2% B27 Supplement (Gibco, 12587010), 1% N2 Supplement (Gibco, 21103049), 1% Glutamine (Gibco, 25030-81), 1% Penicillin/Streptomycin (Sigma-Aldrich, P4333), and 1μg/mL Heparin (Stemcell Technologies, 07980). All cell lines were grown as spheroids and media supplemented with 0.2μg/mL each Human Epidermal Growth Factor Protein (Sino Biological, 10605-HNAE-1) and Human Recombinant bFGF/FGF2 Protein (Sino Biological, 10014-HNAE-1). HEK293T were obtained from American Type Culture collection and maintained in DMEM (Gibco) with L-glutamine (Fisher Sci.), 10% fetal bovine serum (Fisher Sci.), 1% L-Glutamine, and 1% Penicillin/Streptomycin (Sigma-Aldrich, P4333). All cells were grown at 37°C and 5% CO_2_.

### Drugs and small molecules

3-Deazuridine (DAU) was obtained from Santa Cruz Biotechnology (394445) or Cayman Chemical Company (23125). JHU395 and JHU083 were a gift from Barbara Slusher (Johns Hopkins). Prexasertib mesylate (206787) and Prexasertib HCl (206989) were procured from MedKoo Biosciences. Brequinar was procured from SelleckChem (S6626) and Lometrexol from Cayman Chemical Company (18049). 6-diazo-5-oxo-L-nor-Leucine (DON) was procured from Cayman Chemical Company (17580). Guanosine (S2439), Adenosine (S1647), Cytidine (S2053), and Uridine (S2029) were procured from SelleckChem. For *in vitro* use, drugs were reconstituted in DMSO (Millipore Sigma, D2438).

### Proliferation measurements

For Cell Titer Glo, cells were treated with serial concentrations of Prexasertib (0-20nM), DAU (0-30nM), JHU083 (0-20µM), JHU395 (0-20µM), Lometrexol (0-180nM), and Brequinar (0-180nM) or indicated combinations, then incubated for 72 hours. Proliferation was assessed with CellTiter-Glo 2.0 (Promega, G9242) according to manufacturer instructions and quantified using a Spectramax iD5. For BrdU assessment, cells (untreated or treated as indicated) were incubated with 10μM BrdU (ThermoFisher, B5002) for 1hr. After incubation, cells were rinsed with 1X PBS (Cytiva, BSS-PBS-1X6), fixed in ice-cold 70% ethanol, and stored at -20°C. Fixed cells were washed with DPBS and incubated in 2N HCl/0.5% Triton X-100 for 20 min. Cells were stained with anti-BrdU FITC antibody (BioLegend, 364104) for 1 hour and analyzed on a BD Accuri C6 and FlowJo (BD Biosciences).

### Cell cycle measurement

For cell cycle measurement, cells were treated as indicated and fixed in ice-cold 70% ethanol after PBS wash. FxCycle PI/RNase Staining Solution (ThermoFisher, F10797) was added to cells as per manufacturer instruction. Analysis was performed on a BD Accuri C6 and cell cycle analysis performed in FlowJo (BD Biosciences).

### Lentiviral shRNA production

Lentiviral shRNA plasmids (U6 promoter) were designed and produced by Vectorbuilder. shRNA sequences were:

shCTPS1: GCTCTCACATTACCTCCAGAA (TRCN0000045349)

shCTPS2: CGCCTCACCAAGGACAATAAT (TRCN0000369478)

HEK293T cells were plated on a 10cm dish (Corning, 877222) until 60-70% confluent. Cells were transfected with 10μg transfer plasmid (VectorBuilder), 9μg psPAX2 (Addgene, 12260) and 3μg pMD2.G (Addgene, 12259) using Xfect Transfection Reagent (Takara, 631318) according to manufacturer instructions. After transfection, virus was collected in Neurobasal media, filtered through a 0.45μm syringe filter (Globe Scientific, SF-PES-4530-S) and stored at -80°C. Viral particles were added to tumorspheres with 8μg/mL polybrene (Millipore Sigma, TR-1003-G) overnight. Forty-eight hours post-infection, puromycin (Millipore Sigma, P4512-1MLX10) was added at 1μg/mL for 14 days to select for infected cells.

### Western immunoblotting

After indicated treatments, cells were lysed in RIPA buffer (150 mM NaCl; 50 mM Tris pH 8.0; 1.0% IGEPAL CA-630; 0.5% sodium deoxycholate; 0.1% SDS) (Thermo Scientific, PI89900) with Halt protease and phosphatase inhibitor (Thermo Scientific, PI78445). For tumor tissue, tissue was homogenized using an electric homogenizer fitted with a disposable pestle in RIPA buffer. Protein was quantified using Pierce 660nm Protein Assay Reagent (Thermo Scientific, 22660) and immunoblotted with standard procedures. Antibodies and concentrations are listed Table S1.

### Immunofluorescence staining

FFPE sections from flank tumors were deparaffinized and rehydrated with Histoclear (HS2001, Fisher) and ethanol according to manufacturer instructions. Antigen retrieval was performed by pressurized retrieval in pH 6.0 citrate buffer (C9999, Millipore Sigma) for 5 min on high pressure in a pressure cooker (Cuisinart CPC-600 6 Quart 1000 Watt Electric Pressure Cooker). Samples were blocked in 0.1% PBS+TX-100 5% donkey serum (S30-M, Millipore Sigma) for 1 hr at RT prior to staining with antibodies overnight at 4C. Secondary Alexa fluor antibodies (donkey anti-488, 555 or 647; Invitrogen) were diluted at 1:500 in blocking buffer for 1 hour at RT. Samples were mounted in Prolong Gold with NucBlue (P36981, Fisher Scientific). All samples were imaged on a Leica DMi8 microscope with Thunder deconvolution software (LAS X, Leica Microsystems) and quantified in ImageJ.

### Synergy analysis

Visualization and analysis of drug combinations were performed using the SynergyFinder web application (version 3.0, https://synergyfinder.fimm.fi) using viability as readout. Outlier detection was on with LL4 curve fitting. Bliss, ZIP, HSA, and Loewe scores and p-values were calculated as described^35^.

### Mouse xenograft experiments

All animal procedures were approved by Institutional Animal Care and Use Committee (IACUC) of Seattle Children’s Research Institute (Protocol #00000675). Nude female mice (Jackson Laboratories, NU:J-002019 or 007850) were procured at 5 weeks of age and allowed to acclimate for at least 1 week prior to cell implantation. Luciferase-expressing cells were made by transduction of single cells with pLenti CMV mCherry-T2A-Luc blast (a gift from N. Vitanza) followed by hand selection of positive tumorsphere colonies.

### Subcutaneous implantation

Patient-derived MB cells were dissociated with Accutase (AT-104, Innovative Cell Technologies), resuspended in Gibco Neurobasal Medium (21103049, Fisher Scientific), and injected into the flank in a 100 μL aliquot containing 1.25 × 10^6^ cells. Mice were randomly segregated into groups when tumors reached 100 mm^3^ volume: vehicle, Prexasertib (10mg/kg subcutaneous twice daily), 3-Deazuridine (75mg/kg intraperitoneal), or combination until health-defined end points. Tumor size was measured with calipers. For growth curve analysis and survival studies, mice in study cohorts were sacrificed when tumors reached a max of 3000 mm^3^ in total volume. The tumors were excised from sacrificed mice and subjected to routine histopathological processing.

### Orthotopic implantation

MB tumorspheres were treated with Accutase and approximately 100,000 mCherry-positive cells injected per mouse in 2μL of sterile PBS (Cytiva, BSS-PBS-1X6). For surgery, mice were given buprenorphine ER for analgesia and anesthetized with a ketamine/xylazine mixture. A 0.9mm burr hole was made in the right cerebellum (-1.0mm AP 1.0mm ML) from lambda and a stereotactic needle dispensed the cells in two 1μL injections (2.5mm and 2.0mm DV) to deliver the cells. To monitor tumor growth, mice were injected with 150 mg/kg of D-luciferin (LUCK-4G, Gold Biotechnologies), and imaged using the IVIS Spectrum Imaging System 15 minutes post-injection.

### *In vivo* drug administration

JHU083 was prepared in sterile PBS without Ca and Mg (Cytiva, BSSPBS1X6) and administered at a concentration of 20mg/kg via oral gavage twice weekly. Prexasertib mesylate (MEDKoo Biosciences, 206787) or prexasertib HCl (MEDKoo Biosciences, 206989) were prepared in 10% SBE-β-CD (MedChem Express, Cat# HY-17031/CS-0731) and dosed at a concentration of 10mg/kg subcutaneously twice daily. 3-Deazuridine (Santa Cruz Biotechnology, 394445) was diluted in corn oil (Millipore Sigma, C8267-500ML) and dosed at a concentration of 75mg/kg IP once daily.

### Dependency Map Analysis

Dependency data (CERES score) from the Broad Institute’s Pediatric Cancer Dependency Map^20^ was accessed from the 23Q4 data deposit (depmap.org).

### Patient gene expression and proteomic analyses

The processed expression of indicated genes were downloaded from the *R2: Genomics Analysis and Visualization Platform* (http://r2.amc.nl) for indicated data sets^3,23,24^. When calculating expression differences between different groups they were considered significantly different when *P* < 0.05, as calculated using the one-way analysis of variance (ANOVA) test included in the R2 online software. Linear regression was used to define R-values and p-values for correlation plots. For significance of differences between primary and metastatic tumors, the p-value was calculated with an unpaired Student *t* test. For survival analyses in Kaplan–Meier plots, we used a 5-year/60-month time point as a cut-off for survival due to the potential risk of including secondary tumors. The *P* value for survival curves is calculated using a log-rank test of reported patient survival data as described in R2 (https://r2.amc.nl). Only *a priori* fixed cut-offs (median or top 1/3 v bottom 2/3) were used to calculate *P* values in the Kaplan–Meyer survival analysis.

Proteomics data was downloaded from the Medulloblastoma Data Explorer (https://medullo.shinyapps.io/archer2018/). One-way ANOVA was used to determine significant differences.

## Supporting information

Suppementary Figures

## Acknowledgements

We thank the University of Washington/Fred Hutchinson Cancer Center/Seattle Children’s Hospital Cancer Consortium for shared resources including Flow Cytometry and Cell Sorting and Research Pathology supported by NCI P30CA015704. We thank Virginia Hoglund and Galen Stewart for histopathology support, Jim Olson for providing PDX tissue and cell lines, Beth Lawlor for reagent support, Karuna Patil and Rajesh Utamanthil for animal husbandry and veterinary support. Research reported in this manuscript was funded by the National Cancer Institute of the National Institutes of Health under Award Numbers K22CA262343 (MKE) and K22CA258953 (SSP), Kate Amato Foundation (MKE), Steven Higgins Brain Tumor Research Fund, Unravel Pediatric Cancer, the Pediatric Brain Tumor Research Fund of Seattle Children’s, the Run of Hope Seattle, Seattle Children’s Guild Association, and UW/FHCC/SCH Cancer Center Support Grant (P30CA015704).

## Author contributions

**Matthew R Hathaway:** Conceptualization, Data curation, Formal analysis, Investigation, Methodology, Validation, Visualization, Writing – original draft, Writing – review & editing. **Katherine E Gadek:** Data curation, Formal analysis, Investigation, Methodology, Validation, Visualization, Software, Writing – original draft, Writing – review & editing. **Hawa L Jagana:** Data curation, Writing – review & editing. **Isabella C Terrones:** Data curation, Formal analysis, Writing – review & editing. **John M Hemenway:** Data curation, Formal analysis, Writing – review & editing. **Aya Miyaki:** Data curation, Writing – review & editing. **Ashmitha Rajendran:** Investigation, Writing – review & editing. **Michael Meechan:** Data curation, Investigation, Writing – review & editing. **Leonel Elena-Sanchez:** Data curation, Investigation, Writing – review & editing. **Nicholas A. Vitanza:** Methodology, Resources, Writing – review & editing. **Barbara S Slusher:** Methodology, Resources, Writing – review & editing. **Siobhan S Pattwell:** Methodology, Resources, Writing – review & editing. **Myron K Evans II:** Conceptualization, Funding acquisition, Methodology, Project administration, Software, Supervision, Validation, Writing – original draft, Writing – review & editing.

## Figure Legends

**Figure S1: Dependency, gene expression and survival data for additional d*e novo* pyrimidine synthesis and salvage pathway genes in G3 MB. a,** Volcano plot comparing CERES scores of G3 MB cell lines (D283, D425, D458, D341) versus other pediatric CNS tumor lines (n=17) in the Dependency Map using two class comparison. **b,** CERES gene dependency scores for pyrimidine synthesis genes in G3 MB cells (n=4) compared to other pediatric CNS tumor lines (n=17) in the DepMap, 23Q4 release. **c**, CERES scores for normal NSCs (CB660 and U5) for *de novo* pyrimidine synthesis genes as well as *TOP2A* and *RPA2* (controls). **d**, CERES gene dependency scores for pyrimidine salvage genes in G3 MB cells (n=4) compared to all pediatric (left, n=111) and peds CNS (right, n=17). **e**, Relative CTPS1 protein expression in G3a (n=8) and G3b (n=6) tumors from(ref). **f**, Heatmap of relative protein expression for other *de novo* pyrimidine synthesis proteins. **g**, Linear regression of log_2_ normalized *MYC* and *de novo* pyrimidine synthesis gene expression from GSE85217 separated by subtype. **h**, Linear regression of log_2_ normalized *CTPS1* and *CCNB1* expression separated by G3 subtype. **i**, Kaplan-Meier survival of *MYC*-high G3 MB patients separated by top 1/3 versus bottom 2/3 expression of indicated genes. Number of patients are shown on inset.

**Figure S2: DAU treatment alters cell cycle in G3 MB. a,** Cell-cycle analysis of Med-411FHTC cells treated with the indicated concentrations of DAU alone or in the presence of cytidine (C) or uridine (U) for 48h (n=3). *, P < 0.05; **, P < 0.01; ***, P < 0.001, Data are means ± SEM.

**Figure S3: MYC drives expression of pyrimidine and purine synthesis genes, but G3 MB cells are insensitive to purine synthesis inhibition. a**, Representative ChIP-sequencing tracks for MYC in human G3 MB cells at *CTPS1* and *CTPS2* promoters. **b,** Expression (transcripts per million, TPM) of *CTPS1* and *CTPS2* in tumor samples from **a**. **c,** Linear regression of log_2_ normalized *MYC* and *de novo* purine synthesis gene expression in G3 tumors. **d**, CERES gene dependency scores for *de novo* purine synthesis genes in G3 MB cells (n=4) compared to peds CNS (n=17) cell lines. **e**, left - CERES scores for *TYMS* in G3 MB (n=4), all peds (n=111), and peds CNS (n=17); right - Linear regression of log_2_ normalized *MYC* and *TYMS* expression in G3 tumors. **f**, Cell proliferation analysis of Med-411FHTC and Med-2112FHTC tumorspheres treated with indicated concentrations of lometrexol (LTX) for 72h (n=3). *, P < 0.05; **, P < 0.01; ***, P < 0.001, Data are means ± SEM.

**Figure S4: Prexasertib alone has limited effect but synergizes with DHODH inhibition.** Cell proliferation analysis of HD-MB03 and Med-411FHTC tumorspheres treated with increasing concentrations of **a,** prexasertib alone or **b,** brequinar in the presence of indicated doses of prexasertib. Data are means ± SEM.

**Figure S5: JHU083+prexasertib combination limits tumor growth in vivo. a,** chemical structures of DON, JHU083, and JHU395. IVIS imaging of mice orthotopically implanted with **b,** HD-MB03 or **c,** Med-411FHTC -luciferase expressing tumor lines treated as indicated. Images are from mid-point of treatment. Monitored weights of animals from **d,** heterotopic model treated with DAU and/or prexasertib and **e,** orthotopic model treated with JHU083 and/or prexasertib.

