## Supplementary material for "CTPS1 inhibition synergizes with replication stress signaling inhibition in *MYC*-amplified Group 3 medulloblastoma": Suppementary Figures

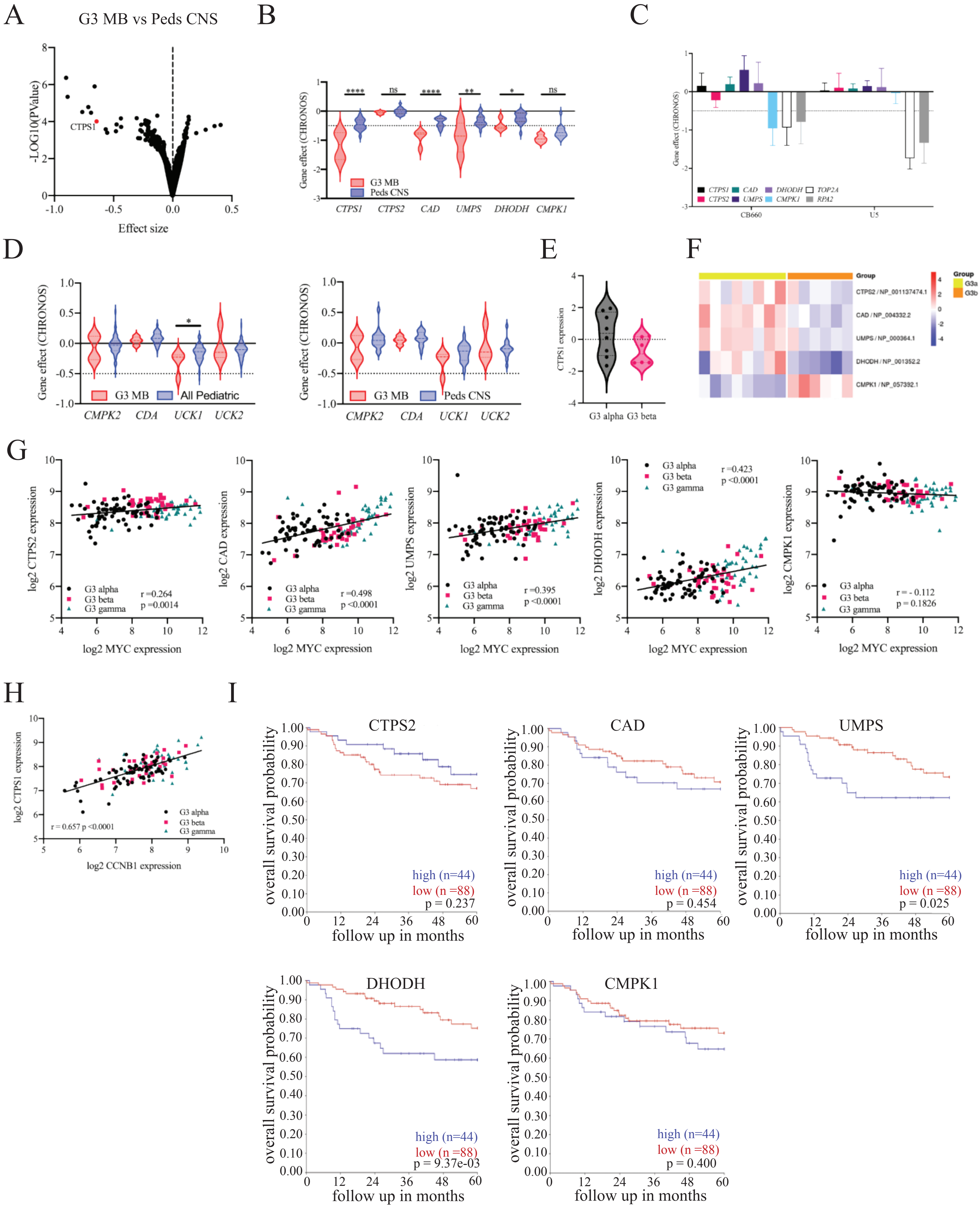


Hathaway et al. Figure S1

A

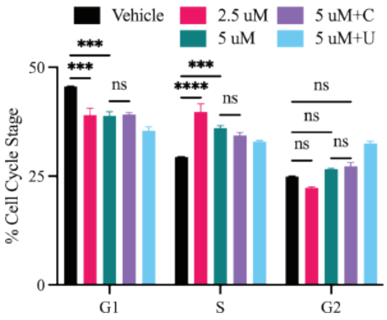


Hathaway et al. Figure S2

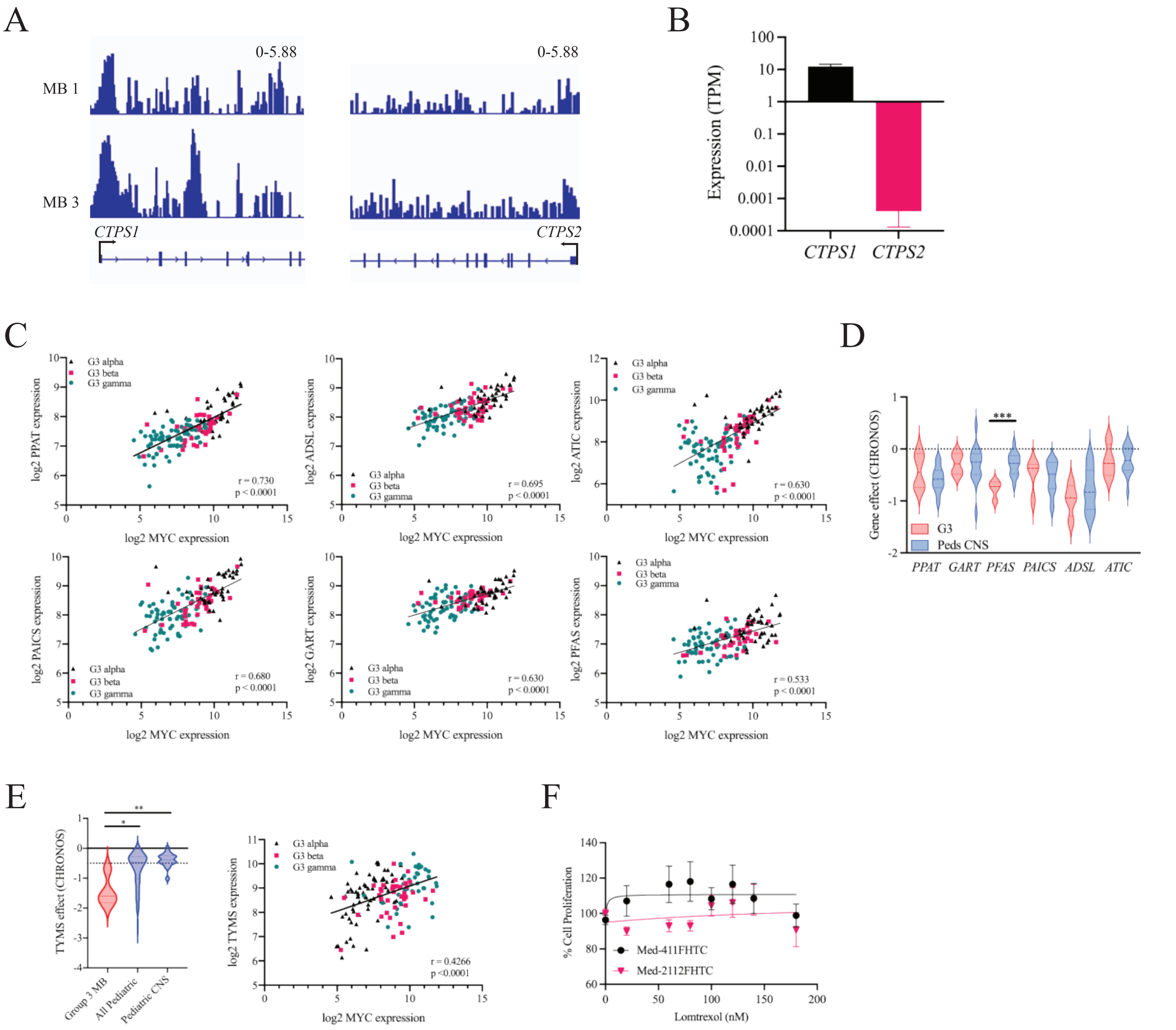


Hathaway et al. Figure S3

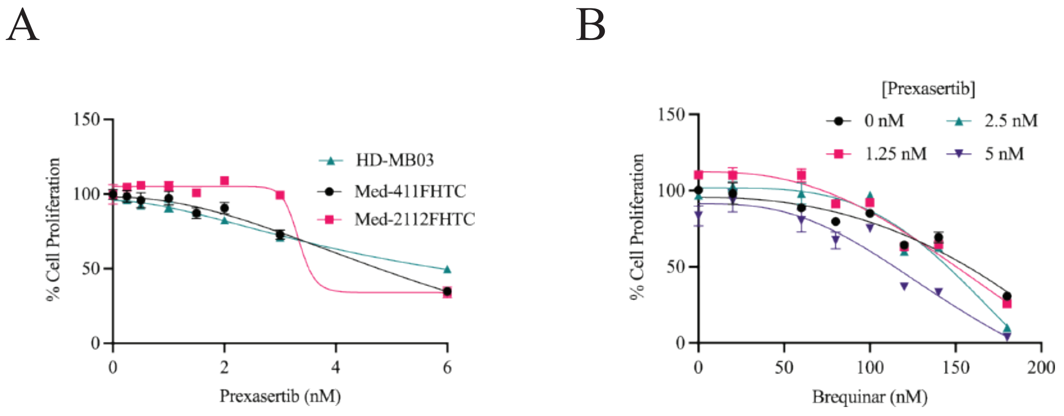


Hathaway et al. Figure S4

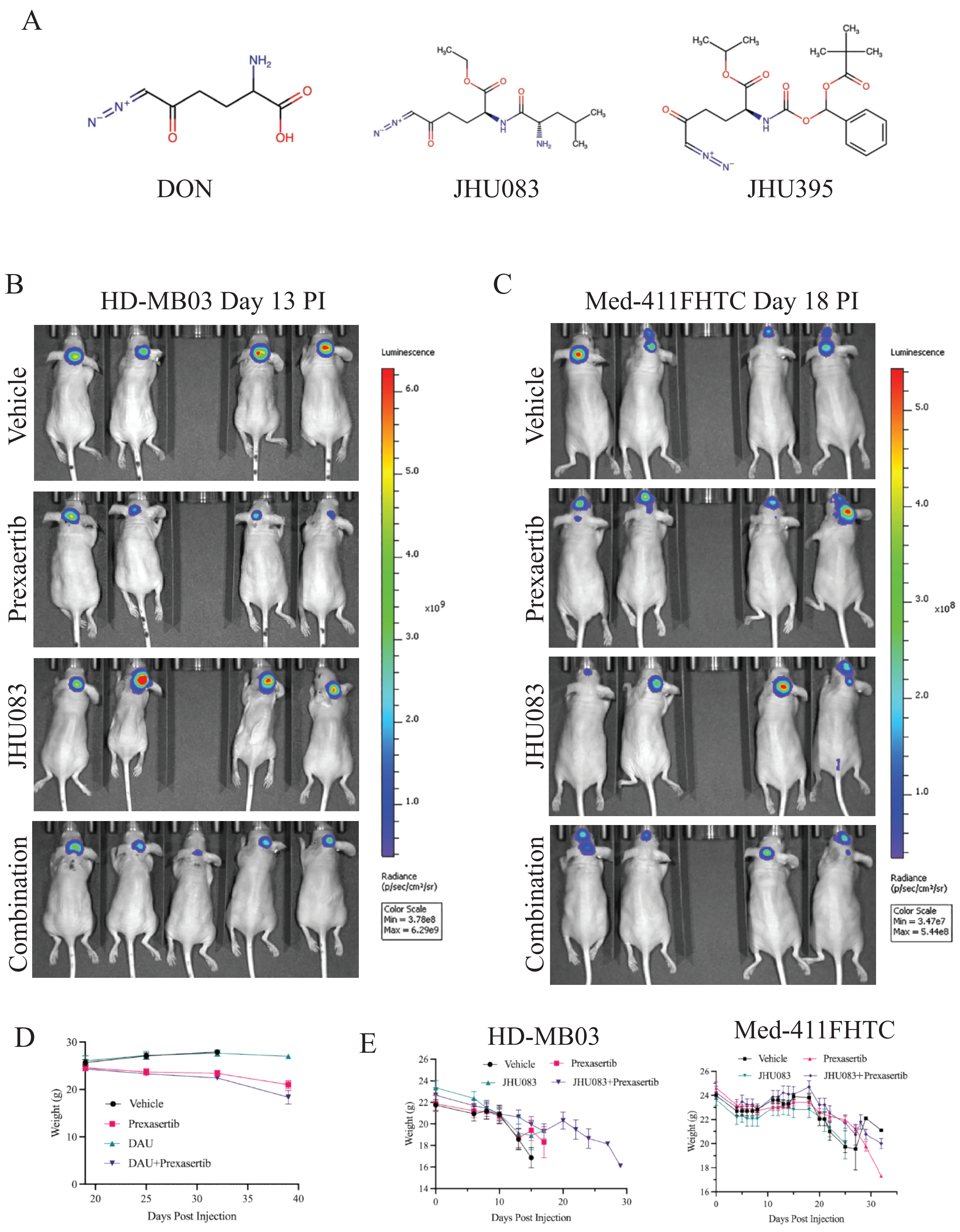


Hathaway et al. Figure S5

Table S1: List of antibodies

| Antibody | Species | Source | Catalogue Number | Dilution |
|---|---|---|---|---|
| Anti-phospho-CHK1 | Rabbit | Cell Signaling Technologies | 2341 | WB (1:1000) |
| Anti-phospho-CHK2 | Rabbit | Cell Signaling Technologies | 2197 | WB (1:1000) |
| Anti-PARP | Rabbit | Cell Signaling Technologies | 9532 | WB (1:1000) |
| Anti-human nuclear antigen | Rabbit | Millipore Sigma | MAB1281 | IHC (1:1000) |
| Anti-CTP Synthase (CTPS1) | Rabbit | Protein Tech | 159141-1-AP | WB (1:1000)<br>IHC (1:200) |
| Anti-pCDC2 | Rabbit | Cell Signaling Technologies | 4539 | WB (1:1000) |
| Anti-pH2AX | Rabbit | Cell Signaling Technologies | 9718 | WB (1:1000)<br>IHC (1:500) |
| Anti-PCNA | Mouse | Cell Signaling Technologies | 2586 | WB (1:1000) |
| Anti-BrdU (FITC conjugated) | Rabbit | Biolegend | 364106 | FACS (4μg/mL) |
| Anti--Actin | Mouse | Sigma Aldrich | A1978 | WB (1:4000) |
| Anti-GAPDH | Rabbit | Millipore Sigma | MAB374 | WB (1:4000) |
| Anti-c-Myc | Mouse | Santa Cruz | SC-40 | WB (1:1000) |
